# Coral niche construction: coral recruitment increases along a coral-built structural complexity gradient

**DOI:** 10.1101/2021.10.14.464352

**Authors:** Viviana Brambilla, Andrew H. Baird, Miguel Barbosa, Inga Dehnert, Joshua T. Madin, Clare Peddie, Maria A. Dornelas

## Abstract

Niche construction is the process through which organisms modify environmental states in ways favourable to their own fitness. Here, we test experimentally whether scleractinian corals can be considered niche constructors. In particular, we demonstrate a positive feedback involved in corals building structures which facilitate recruitment. Coral larval recruitment is a key process for coral reef persistence. Larvae require low flow conditions to settle from the plankton, and hence the presence of colony structures that can break the flow is expected to facilitate coral recruitment. Here, we show an increase in settler presence on artificial tiles deployed in the field along a gradient of coral-built structural complexity. Structural complexity had a positive effect on settlement, with an increase of 15,71% of settler presence probability along the range of structural complexity considered. This result provides evidence that coral built structural complexity creates conditions that facilitate coral settlement, while demonstrating that corals meet the criteria for ecological niche construction.

## 1. INTRODUCTION

Niche construction is the process by which organisms modify their surrounding environment in ways that may affect their evolution, and/or the evolution of other organisms that experience the modified conditions (Matthews et al. 2014, Laland et al. 2016). Niche constructor organisms must meet three nested criteria, the first two characterizing niche construction, and the third determining whether niche construction generates an evolutionary response (Matthews et al. 2014). First, the niche constructor must change its external environment, through behavioural, physical/chemical or other metabolic processes (Donohue 2014, Laland et al. 2016). Second, these modifications must bias natural selection upon the organism itself and/or other organisms, either positively or negatively (Zahavi 1974, Matthews et al. 2014). Third, the modifications must leave a trace in the evolutionary history of the organisms involved, in the form of an evolutionary response to the environmental modification. (Matthews et al. 2014). While criteria 1 and 2 can be tested in an ecological framework, criterion 3 applies to an evolutionary time scale. The first two criteria describe an ecological feedback loop that can lead to diverse consequences, ranging from the local extinction of the responding population, to triggering trait fixation (criterion 3).

Ecosystem engineering species are a class of putative niche constructors (Matthews et al. 2014, Laland et al. 2016) since by producing long-term environmental changes they can affect macroevolutionary patterns and biodiversity (Erwin 2008). Scleractinian corals, as autogenic bioengineers (Jones et al. 1994), are a prime example of ecosystem engineering since they physically create reefs that harbour some of the most biodiverse communities in the world (Hughes et al. 2017). Yet, there are no explicit tests of their niche construction capability. Here, we focus on the effects of skeletal three-dimensional structures on corals themselves, examining whether this trait facilitates recruitment, thereby establishing a positive ecological feedback. We aim to advance understanding of coral niche construction by testing whether corals meet the second criterion of niche construction (Fig.1).

**Figure 1.**
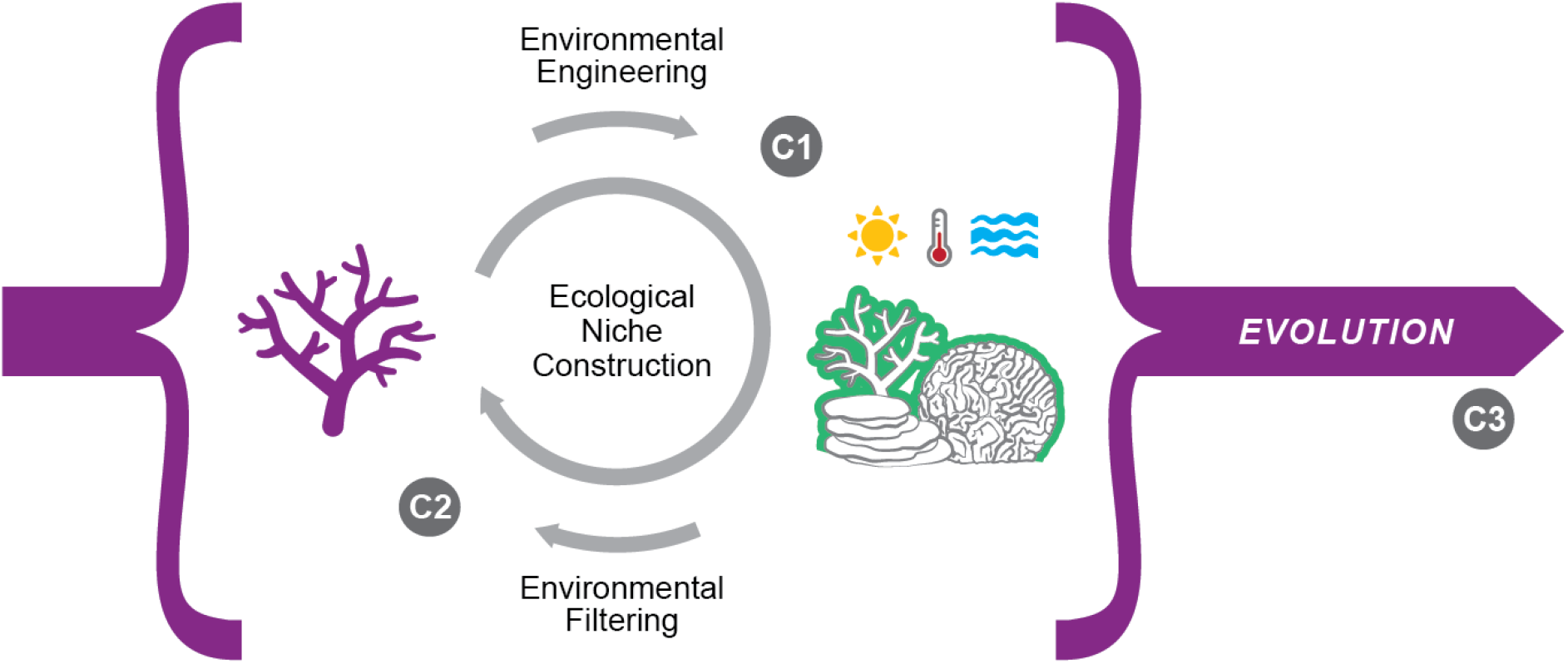
Evolution of coral through niche construction. Ecological processes such us ecosystem engineering and environmental filtering, both detectable at ecological time scales, plausibly play an important role in coral evolution. Corals are obligate physical ecosystem engineers, since they create and modify the habitat around themselves. Their physical structures inherently modify the environmental conditions that the colonies themselves will experience. Transforming the 3D structure of the reef is likely to bring changes in environmental patterns (flow, light and temperature), as well as creating habitat and resources that other marine species exploit and thereby impact community composition, which in turn plausibly changes the selective pressures on the coral. Over time, these ecological processes are likely to shape coral evolution. In the grey circles, the three criteria for niche constructions outlined in the introduction are paired to the presented processes. C1 = criterion 1, C2 = criterion 2, C3 = criterion 3.

Scleractinian corals build complex and heterogeneous environments, which harbour some of the most biodiverse and threatened communities in the world (Hughes et al. 2017). Corals have a planktonic life stage and larval recruitment success is key to the persistence of the reef ecosystem (Bellwood et al. 2004). At the end of the planktonic stage, coral larvae need to settle on suitable substratum, metamorphose and start the benthic life. As with other benthic marine organisms, corals undergo severe early-life stage bottlenecks, and recruitment success depends on both abiotic and biotic factors (Ritson-Williams et al. 2009). For example, crustose coralline algae (CCA) release chemical cues that induce the coral to settle and metamorphose (Heyward & Negri 1999). In contrast, macroalgae can compete with coral for space occupancy and negatively affect coral recruitment (Mumby et al. 2006). Once metamorphosized, post-settlement processes transform a settled polyp into a coral colony, which through its hard skeleton modifies the topography of the reef. Coral colonies modify the overall complexity of reef habitats both when alive and after death, when they leave behind their hard skeleton (Richardson et al. 2017) as ecological inheritance (Odling-Smee et al. 2013). Reef structural complexity is a measure of how corals engineer the environment, and is important for ecosystem function and maintenance from an ecological perspective (Graham & Nash 2013, Zawada et al. 2019b). For example, reef structural complexity provides microhabitats and determines fish assemblage structure (Nash et al. 2014). Furthermore, heterogeneity in coral colonies change the local environmental conditions, such as light (Brakel 1979) and water flow (Hench & Rosman 2013) to create a range of microhabitats and niches. All this demonstrates that corals meet the first niche construction criterion suggesting that coral niche construction may might be one process influencing the evolution of this diverse and productive ecosystem (Laland et al. 2015).

Moving to the second criterion, positive ecological feedbacks to niche-constructing populations have been identified in other organisms (Matthews et al. 2014), but not in corals. In the tundra, for example, plant species of all growth forms (i.e. forbs, grasses, sedges, deciduous shrubs and evergreen shrubs) collectively modify niches independently of local environmental conditions, increasing taxonomic diversity (Bråthen & Ravolainen 2015). These environmental modifications bias natural selection upon the niche constructor and other associated species with important evolutionary consequences (Laland et al. 2015). Another example is the sediment bioturbation and thickness of shell beds in paleoecological records, which increased over geological time as result of increased ability of the organisms to modify ecosystems (Erwin 2008). This process resulted in greater evolutionary diversification of benthic niche constructors and ecosystem engineers as well (Erwin 2008). In modern coral reefs, we can focus on how corals increase their own fitness modifying the environment in predictable and favourable ways. Identifying coral traits that capture these modifications and feed back to coral fitness would allow us to show that the second criterion for coral niche construction is met.

Settlement success plays a key role in coral fitness, because this is the life stage with lowest success rate. There is evidence that millimetre-scale rugosity of the substratum enhances settlement success (Birkeland & Randall 1981, Hata et al. 2017). However, the extent to which coral settlement is affected by increased habitat complexity (at the centimetre to meter scale) built by coral colonies remains unclear. Coral larvae are poor swimmers, and often rely on eddies created by small structural obstacles, such as sea urchin burrows in the field (Birkeland & Randall 1981) or 1-cm blocks in the lab (Hata et al. 2017), to be able to find suitable substratum and attach. On coral reefs, these flow conditions can be built by adult coral colonies with different structural complexity (Zawada et al. 2019a b). Areas of flow recirculation and of reduced current created by the presence of structural 3D complex coral assemblages (Hench & Rosman 2013, Zawada et al. 2019a) can play an important role in creating ideal hydrodynamic conditions for settlement and attachment of the larvae. Thus, high structural complexity built by corals is predicted to be favourable for coral settlement and recruitment.

Here, we show that assemblages of higher coral 3D complexity structures have higher probability of settlement of coral larvae. Specifically, we measured settlement on tiles used to hold either dead or alive corals during a reciprocal transplant experiment. We predicted that tiles in more complex coral assemblages (coral-built structural environments) will have greater probability of having coral settlers than tiles in less complex assemblages. Dead skeleton persisting over generational time as ecological inheritance can affect the evolutionary trajectories of the niche constructor as well. As such, we further investigate the effect of the status of the coral on tile (alive or dead) on recruitment success.

## 2. MATERIALS & METHODS

A coral reciprocal transplant experiment was set up in the South-East lagoon of Maghoodoo Island (3°04′N, 72°57′E, Republic of Maldives) in January 2017 (Fig.2). The experiment used 370, ~10 cm long fragments of colonies belonging to four species of branching corals commonly found in the lagoon (*Acropora divaricata*, *A. muricata, Porites rus*, *P. cylindrica*). Each fragment was cemented with reef cement (NYOS © reef cement) to a concrete disk tile (7×2,5cm) and then attached to one of 10 racks in either a shallow (5 racks, 5-6 m) or a deep (5 racks, 16-18m) site. At the start of the experiment, each rack had 37 concrete tiles with same-size same-species living fragments attached (Fig.2). All the racks had similar structural complexity in January 2017, but each one of them was left in different experimental conditions (deep site = low light, shallow site = high light) for 15 months (Fig.2). As a result, coral growth rate and mortality were different among racks, reflecting different environmental conditions experienced by the coral fragments. By the end of the experiment (May 2018), each rack had a different number of living fragments that grew into colonies (Fig.2), while the dead ones had different sizes and shapes depending on the time of death. Thus, the complexity of each rack increased during the experiment, but each rack displayed a different degree of structural complexity at the end.

**Figure 2.**
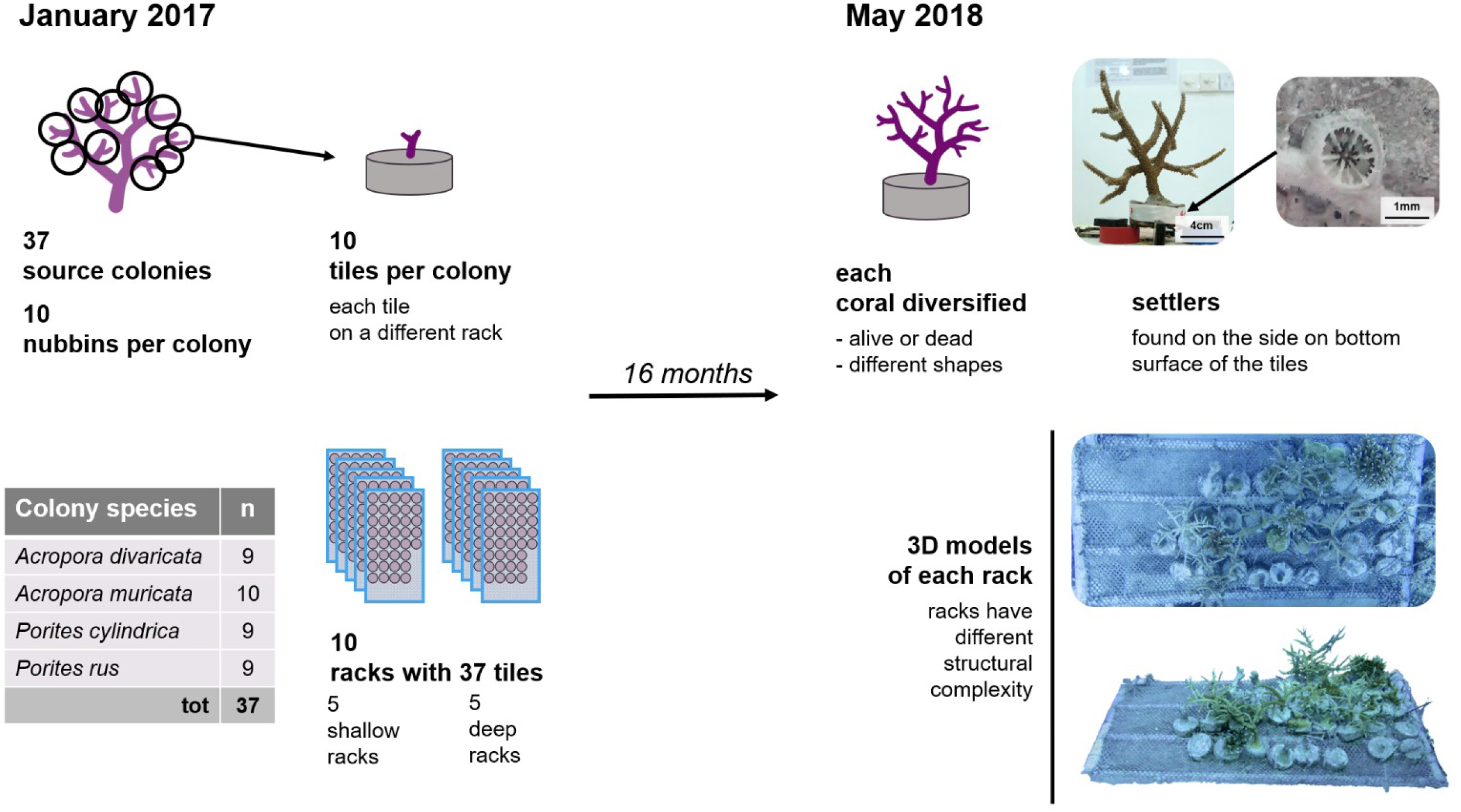
**Timeline of experimental setup**, from January 2017 to May 2018.

**Figure 3.**
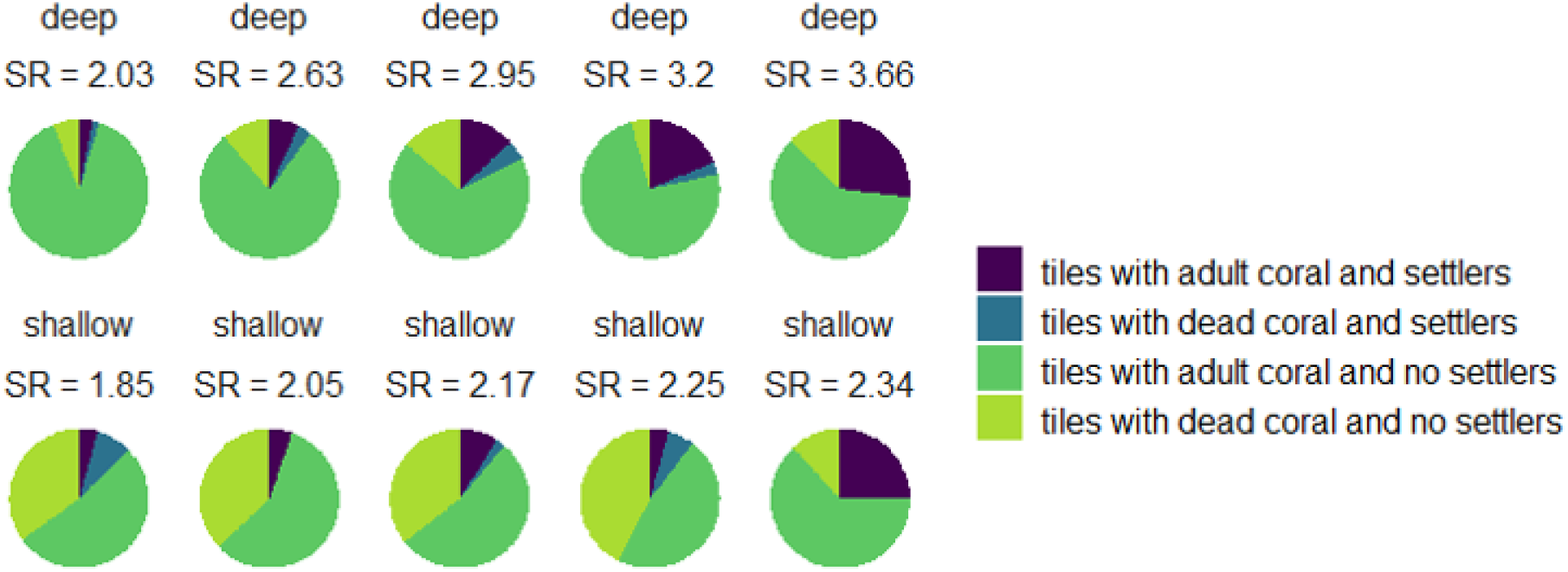
Tile proportion in each rack. Proportion of tiles with live corals and settlers (dark blue), tiles live corals but without settlers (light green), tiles with dead coral and settlers (ocean blue) and tiles with dead corals and no settlers (lettuce green) Racks are arranged by increasing SR at each depth.

A significant coral spawning event was observed in the Maldives on April 1^st^ 2017 (ID, personal observation). The coral larvae pool was expected to be approximately the same for all the racks, since they were in the same enclosed lagoon. Given the coral assemblage present on the island (Montano et al. 2012), we predicted that more than just the species used in the experiment were spawning simultaneously in the lagoon. All the racks were retrieved and brought in the lab for tile analysis in the second week of May 2018 (6^th^ and the 11^th^ of May 2018). Settlers were found on the bottom and lateral surface of the concrete tiles used for the transplant experiment (Fig.2). Crustose coralline algae (CCA), bryozoans, molluscs, and sponges were also observed on the tiles, but cover was not quantified. The status (alive or dead) of the fragment attached to every tile was recorded. 26 to 37 tiles per rack were analysed, corresponding to a total of 214 tiles with a live coral attached, and 125 with a dead coral attached. Tiles were bleached overnight in a 10% commercial bleach solution, rinsed and dried, and then examined under microscope as per [15]. Bigger settlers (> 2 mm) were considered juvenile corals that settled in 2017 and left out of the analysis (Babcock et al. 2003). Smaller settlers (< 2mm, Fig.2) provided an estimate of settlers from the larval supply of 2018 (Babcock et al. 2003) and were used as a measure of fitness (Hunt & Hodgson 2010).

Given the slow growth rate of corals it is plausible to assume that the rugosity of the racks did not change significantly in the last month of underwater permanence. We used surface rugosity (SR) of the rack at the end of the experiment as a measure of local structural complexity at time of settlement, which occurred less a month before we retrieved the racks. SR was estimated as the ratio between the 3D surface area and its planar orthogonal projection of the surface on the horizontal plane (Friedman et al. 2012), with values that range from 1 (perfectly flat surface) to infinity. 3D digital models of each rack surface (Fig 2) were obtained through structure-from-motion photogrammetric techniques (Westoby et al. 2012). A total of 160 to 190 pictures were taken of each rack from different angles underwater (House et al. 2018) with a Powershoot camera (Canon 5X). Pictures were then analysed in Agisoft Photoscan Professional (Agisoft LLC 2018). The surface area of each rack was computed with the built-in function and used to calculate SR. The values of SR were mean standardized for analysis.

We fitted binomial Bayesian generalized linear mixed models to examine the effect of SR on explaining the presence of settlers. To control for the effect of the status of the fragment attached to the tile (i.e. alive or dead) and the depth of the rack (i.e. shallow or deep), we fitted a total of 7 models: a model including SR, depth and the status of the fragment; 3 models with any combination of 2 of the variables, and 3 models including the effects of one variable at the time. Rack ID was included in all models as random effect to account for unexplained environmental differences between experimental racks. All the priors were left as default values, and for each model four chains for 20000 iterations were run, with a warm-up period of 1000 iterations and a thinning rate of 10 iterations. We ensured that R^ values were almost 1 and goodness of fit was assessed by visual inspection of the chains. We used the Widely Applicable Information Criterion (WAIC) to evaluate the relative goodness of the fit (Vehtari et al. 2017), and check consistency of best fit with the leave-one-out cross-validation (LOOic) (Vehtari, A., Gelman, A., and Gabry 2016). All analyses were performed in R (version 3.3.2, (R Core Team 2018)). Models were fitted with the probabilistic language RStan using the ‘brms’ (Bürkner 2017) and ‘loo’ (Vehtari, A., Gelman, A., and Gabry 2016) packages.

## 3. RESULTS & DISCUSSION

Racks surface rugosity (SR) ranged from 1,85 to 3,66. Some corals died before the end of the experiment, and some of these had begun to erode. Therefore, as expected, lower levels of coral fragment mortality led to more structurally complex racks. Values of SR were well distributed along the range, which is representative of a healthy Indo-Pacific reef. Similar SR values were found on Australian reefs (Bryson et al. 2017) and the presence of grown branching corals with convoluted shapes led to higher SR values on the most complex racks (Figueira et al. 2015).

Settlers were found on 50 out of 339 tiles. We consistently detect an effect of SR, regardless depth or presence of live coral on the same tile (Table 1). Models that included rack SR had lower WAIC and LOOic (Table 1). Moreover, the model with SR as the only predictor variable (Fig.4 a-b) had the best goodness of fit according to both criteria (Table 1). In this model, the estimated effect size of SR on settlement probability was 0,38, with credible intervals not overlapping zero (Fig.4 b). This corresponds to an increase of 15,71 % of settler presence probability along the range of complexity considered. Together our results show that the probability of settlement increased with local structural complexity and highlight the importance of coral-generated structures for the beginning of benthic life stage of these organisms. Greater micro-scale complexity of the substratum enhances coral larval settlement (Hata et al. 2017), our study provides evidence that larger scale coral-built habitat complexity has a positive effect on coral settlement as well, enhancing coral fitness.

**Table 1.** Models results. Effect sizes of the variables, Widely Applicable Information Criterion (WAIC) and leave-one-out cross validation information criteria (LOOic) for all the models. In grey, variables whose 95% Credible Interval (CrI) do not overlap with 0. Models are arranged by increasing WAIC and LOOic. All the models included rack ID as random factor.

| Model | Variable | Estimate | lower<br>95% CrI | upper<br>95% CrI | WAIC<br>(se) | LOOic<br>(se) |
| --- | --- | --- | --- | --- | --- | --- |
| presence ~<br>SR | intercept | -1,84 | -2,26 | -1,47 | 283,38<br>(23,41) | 283,54<br>(23,50) |
|  | SR | 0,38 | 0,02 | 0,78 |  |  |
| presence ~<br>SR + site | intercept | -2,13 | -2,94 | -1,43 | 284,47<br>(23,70) | 284,58<br>(23,72) |
|  | SR | 0,57 | 0,02 | 1,19 |  |  |
|  | site: shallow | 0,54 | -0,68 | 1,76 |  |  |
| presence ~<br>SR + status | intercept | -1,82 | -2,31 | -1,39 | 285,45<br>(23,72) | 285,43<br>(23,72) |
|  | SR | 0,37 | 0,01 | 0,77 |  |  |
|  | status: dead | -0,09 | -0,78 | 0,59 |  |  |
| presence ~<br>status +<br>SR + site | intercept | -2,11 | -2,92 | -1,39 | 286,53<br>(23,98) | 286,55<br>(23,99) |
|  | status: dead | -0,13 | -0,83 | 0,54 |  |  |
|  | SR | 0,57 | 0,03 | 1,17 |  |  |
|  | site: shallow | 0,57 | -0,63 | 1,85 |  |  |
| presence ~<br>site | intercept | -1,68 | -2,38 | -1,07 | 287,24<br>(23,56) | 287,46<br>(23,58) |
|  | site: shallow | -0,34 | -1,31 | 0,58 |  |  |
| presence ~<br>status | intercept | -1,76 | -2,31 | -1,29 | 288,06<br>(23,63) | 288,15<br>(23,64) |
|  | status: dead | -0,21 | -0,90 | 0,46 |  |  |
| presence ~<br>status +<br>site + | intercept | -1,65 | -2,30 | -1,05 | 289,30<br>(23,82) | 289,51<br>(23,90) |
|  | status: dead | -0,13 | -0,85 | 0,56 |  |  |
|  | site: shallow | -0,31 | -1,27 | 0,61 |  |  |

**Figure 4.**
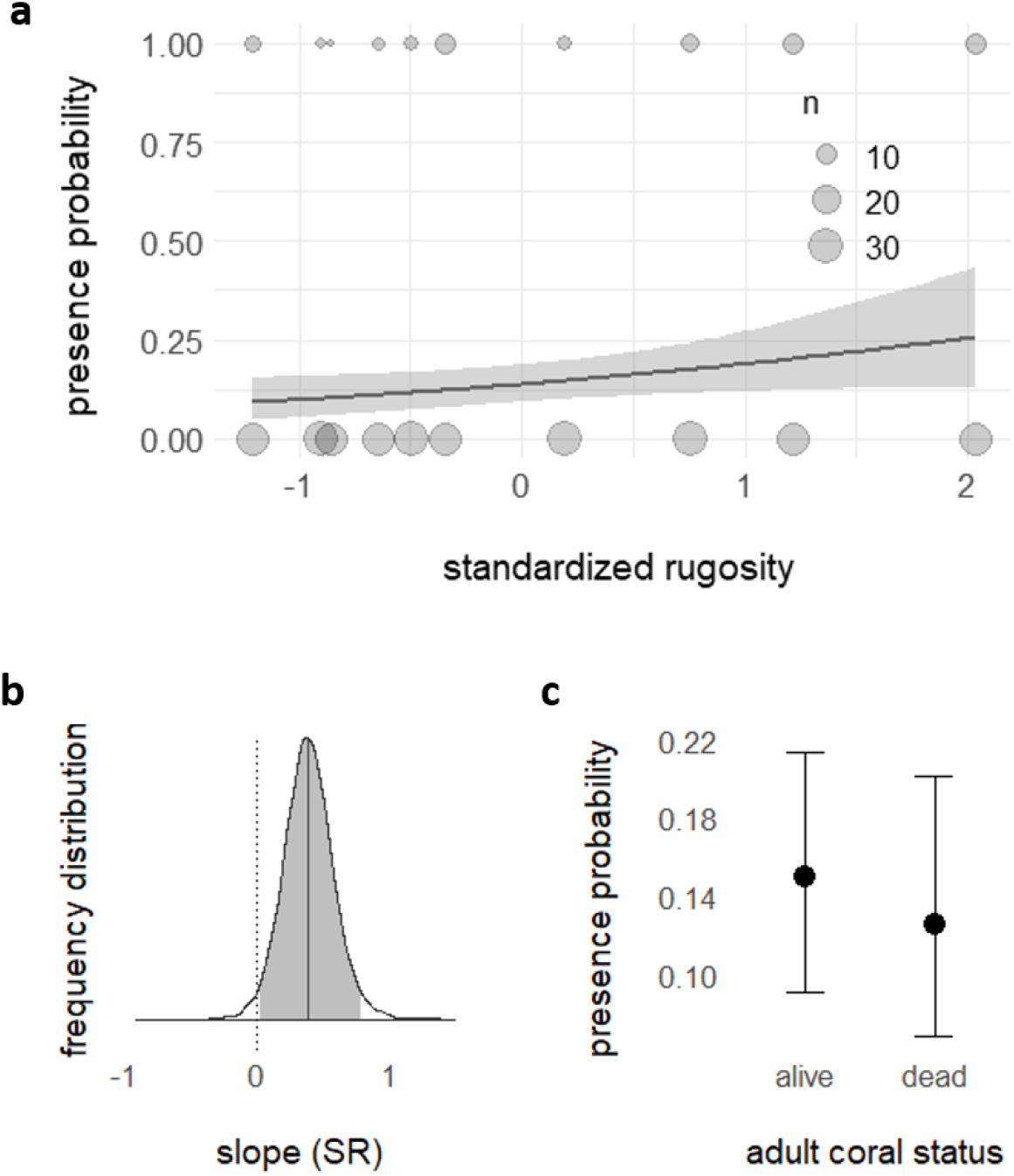
Relevant models results. a) Marginal effect plot for the best fit model, which included only surface rugosity (SR) as fixed effect. Dots represent the number of tiles where settlers and juveniles were found or not. b) Posterior distribution of the slope of the best fit model with 95% credible intervals. c) marginal effect plot for the model which included the status of the adult coral attached to the tile as only explanatory variables, of which we did not detect an effect.

We failed to detect and effect of status of the coral attached to the tile on settling probability (Table 1, Fig.4 c). This suggests that there was probably no biotic interaction between adults and settlers on sides and bottom surfaces of the tiles. As predicted, the presence of corals itself did not have an effect, but more importantly the complexity of their forms determined the positive ecological feedback that we found. Higher complexity has higher chances of diminishing water flow and creating eddies that lead the larvae towards the bottom (Zawada et al. 2019b). Skeleton structures from previous generations of corals can be considered as ‘ecological inheritance’ (Odling-Smee et al. 2003) regardless of colony survival. An ecosystem engineer leaves ecological inheritance when the modification of the environment persists longer than the life-time of the ecosystem engineer (Odling-Smee et al. 2003). Here, this modification (i.e. structural complexity of the skeleton) potentiates coral fitness intergenerationally, and ecological inheritance contributes to the niche construction process.

Although light is a major factor affecting coral growth (Buddemeier & Kinzie 1976, Hoogenboom & Connolly 2009) and zonation (Wellington 1982), we failed to detect an effect of light on settlement. Effects of light would likely have been more detectable when considering recruitment on tile topsides (Vermeij 2006), which did not occur during our experiment. Furthermore, coral structures can also shadow the benthos, modulating the effect of light on the bottom surface (Brakel 1979). Here, we measured fitness as settlement success (Hunt & Hodgson 2010). Light may play a more important role in later coral ontogeny and certainly the effect of structural complexity on settler survivorship needs further investigation. Further experiments including non-coral-built structures and zero-complexity structures as controls could elucidate about the role of coral-built structures in enhancing coral fitness when compared to natural conditions. Nonetheless, our findings provide strong evidence that along a gradient of increasing structural complexity, settler presence increases as well.

As a metaphor, environmental filtering has been used to describe specific values of abiotic environmental variables that “filter out” certain species or traits, not allowing them to persist in specific areas (Keddy 1992, Kraft et al. 2015). In coral reefs, the environmental filtering concept can explain some aspects of reef zonation. For example flow conditions can filter out morphs not suited to face specific hydrodynamic forces and leave structurally clustered species coexisting under similar flow regimes (Madin & Connolly 2006). The concept of environmental filtering (Keddy 1992) has recently been criticized for being used incorrectly (Kraft et al. 2015, Cadotte & Tucker 2017, Thakur & Wright 2017), especially when considering ecosystem engineers (Thakur & Wright 2017). Problems arise when inferring the environmental filter from species or trait observational data, since environmental gradients can simultaneously affect other coexistence mechanisms, like competition for resources, or bioengineer activity (Kraft et al. 2015, Cadotte & Tucker 2017). Niche construction can sustain micro-modifications of the local environment at a small (individual) scale, affecting the local community interactively with the macro-environment (Cadotte & Tucker 2017, Thakur & Wright 2017). The latter seems to be the case for corals, since the differences in growth rates and forms of corals caused by environmental conditions promote heterogeneity in ecosystem functions (Zawada et al. 2019b). This results in an increase in the heterogeneity of community assemblages that in turn shape the overall environment in multitude ways. The environment cannot be considered independently, since reef habitats are literally built by their foundational organisms. The findings of our experiment imply that different recruitment rates resulted from the dynamic construction of different microenvironments (as a consequence of the presence of different coral colonies) within the same macro-environment (two sites in the same lagoon). Since dependent on engineering activity, recruitment and new coral occurrence cannot be explained by biotic, abiotic or dispersal local conditions separately, but rather by a positive feedback interaction of all of the above. Focusing on environmental filtering overlooks this intricate network of reciprocal causation between corals and the environment.

Coral facilitation of settlement has also implications for recovery from disturbances. Human induced disturbances to coral reefs, such as temperature and acidification, are predicted to increase (Hughes et al. 2017), together with the scale of the impacts on the reefs (Hughes et al. 2003). Nonetheless, reef recovery from mass mortality is possible and coral larvae settlement is a necessary process for this recovery. Our findings offer a mechanistic explanation for increased rates of recovery at sites with higher levels of structural complexity due to coral presence (Graham & Nash 2013). Nevertheless, rigorous experimental tests and a better understanding of the mechanisms underlying coral niche construction is urgent and timely in order to promote ecosystem post-disturbance recovery. Indeed, by regulating habitats at a local scale, ecosystem engineering and niche construction can establish population and ecosystem feedbacks and maintain ecosystem health and resilience (Boogert et al. 2006). The experimental system developed here, which can flexibly manipulate the composition, structure and species identity of coral pieces on racks, offers considerable potential to explore these issues further.

The ecological and evolutionary implications of our findings deserve attention. In forest ecology, wildfires, which were considered a purely extrinsic factor, have been shown to be dependent on a set of niche-constructing flammability traits (e.g. branch-morphologies, self-pruning ability, leaf-size, oil content) (Schwilk 2003, Schwilk & Caprio 2011). This demonstrates how organism features can influence external environment in ways that modify selective feedback and eventually their evolution (Schwilk 2003, Post & Palkovacs 2009, Schwilk & Caprio 2011). Coral shapes have an important role in shaping the evolutionary history of other taxa. For example, the emergence of coral branching morphologies are key for the diversification of small-size fish (Bellwood et al. 2017). Yet, whether morphological coral traits affect the evolutionary history of coral groups remained undetermined. Given the three criteria for niche construction (Matthews et al. 2014), we now show that corals meet criterion 2: they modify selection pressure upon themselves through modification of the environment. Defining traits that enhance population fitness enables to look at the evolutionary history of such traits, creating the ground for the test of criterion 3, i.e. studying the evolutionary history and phylogeny of such traits. Models that can predict a range of coral complexity traits from size and species are becoming available in the literature (House et al. 2018, Zawada et al. 2019b a). This allows the use of geological datasets to investigate the evolution and prevalence of 3D traits of interest and the role of niche construction in coral evolution. Here, we make the first step forward in defining coral niche construction, presenting structural complexity as a niche-constructing trait in coral reef ecosystems.

## Acknowledgements

This work was funded by the School of Biology of the University of St Andrews and the Templeton Foundation (grant #60501, ‘Putting the Extended Evolutionary Synthesis to the Test’). MB was supported by a postdoctoral fellowship from Fundação para a Ciência e a Tecnologia (SFRH/BPD/82259/2011). The funders had no role in study design, data collection and analysis, decision to publish, or preparation of the manuscript. The research approved by the Ministry of Fisheries and Agriculture of Maldives (protocol numbers: (OTHR)30-D/INDIV/2016/537 and (OTHR)30-D/lNDlV /2078/739). We thank Kevin Laland for comments and feedback on the niche construction angle, and Fabio Guzzo for help in producing conceptual figures. We thank MaRHE center for support in the field and the Behaviour and Biodiversity group at University of St Andrews, especially Laura Antão for feedback and suggestions during the data analysis.

